# CHCHD2 mediates glioblastoma cell proliferation, mitochondrial metabolism, hypoxia-induced invasion, and therapeutic resistance

**DOI:** 10.1101/2022.07.05.498855

**Authors:** Jan C. Lumibao, Payton Haak, Vladimir L. Kolossov, Jee-Wei Emily Chen, Jeremy Stutchman, Alejandra Ruiz, Mayandi Sivaguru, Jann N. Sarkaria, Brendan A. C. Harley, Andrew J. Steelman, H. Rex Gaskins

## Abstract

Glioblastoma (GBM) is the most common and malignant primary brain tumor in adults and remains incurable. The mitochondrial coiled-coil-helix-coiled-coil-helix domain-containing protein 2 (CHCHD2) is demonstrated to mediate mitochondrial respiration, nuclear gene expression, and cell migration, but evidence of this in GBM is lacking. We hypothesized that CHCHD2 would serve a functional role in U87 GBM cells expressing the constitutively active epidermal growth factor receptor variant III (EGFRvIII). Amplification of the *CHCHD2* gene was found to be associated with decreased patient overall survival and progression-free survival. CHCHD2 mRNA levels were increased in high-versus low-grade glioma, *IDH*-wt GBMs, and in tumor versus non-tumor tissue. Additionally, CHCHD2 protein expression was greatest in invasive, EGFRvIII-expressing patient-derived samples. CRISPR-Cas9-mediated knockout of CHCHD2 in EGFRvIII-expressing U87 cells resulted in altered mitochondrial respiration and glutathione status, decreased cell growth and invasion in both normoxia and hypoxia, and increased sensitivity to cytotoxic agents. CHCHD2 was distributed in both mitochondria and nuclei of U87 and U87vIII cells, and U87vIII displayed greater nuclear CHCHD2 compared to isogenic U87 cells. Incubation in hypoxia, serum starvation, and reductive unfolding of CHCHD2 induced nuclear accumulation of CHCHD2 in both cell lines. Collectively, these data indicate that CHCHD2 mediates a variety of GBM cell hallmark characteristics and highlights mitonuclear retrograde signaling as a pathway of interest in GBM cell biology.

**Implications:** These data demonstrate CHCHD2 as a mediator of a number of GBM cell functions representing disease hallmarks, as well as highlight its subcellular distribution in response to metabolic stressors. These results may inspire therapeutic strategies undermining mitochondrial biology to potentially improve GBM tumor management.

## Introduction

Glioblastoma (GBM, WHO grade IV glioma) is the most common, malignant, and aggressive form of primary brain tumor in adults, accounting for approximately 50% of diagnosed gliomas each year [1]. Patients with GBM present with a median survival time of only 15-20 months, with only 5-10% of patients surviving after 5 years [2]. Despite the current multimodal standard of care, which consists of maximal surgical resection followed by radiotherapy and chemotherapy with the DNA alkylating agent temozolomide (TMZ), overall prognosis remains poor, underscoring the need for a deeper understanding of tumor biology to inspire new therapeutic targets. Contributing to tumor aggressiveness and virtually universal recurrence are GBM hallmarks, including but not limited to: rapid, diffuse invasion into surrounding brain parenchyma, substantial chemo-and radioresistance, and rapid adaptation to microenvironmental stressors such as hypoxia [3, 4].

Amplification of the epidermal growth factor receptor (*EGFR*), one of the most common genetic abnormalities observed in GBM tumors [5], leads to increased proliferative and anti-apoptotic signaling, as well as invasive behavior [6, 7]. Tumors that exhibit *EGFR* amplification also frequently present with the constitutively active EGFR variant 3 (EGFRvIII) mutant [5], which arises from deletion of exons 2-7 of the *EGFR* gene and results in a truncated, yet constitutively active, EGFR protein [6]. The resultant increased signaling downstream confers enhanced glioma malignancy through multiple mechanisms, and, importantly, the EGFRvIII mutant is not present on non-malignant tissues [6]. However, although targeting EGFRvIII is a rational strategy to combat GBM, phase II clinical trials with the EGFR receptor tyrosine kinase inhibitor erlotinib and phase III clinical trials with the EGFRvIII vaccine rindopepimut have failed to robustly increase patient overall survival, highlighting the immense plasticity of GBM cell populations, which dampens treatment efficacies and hinders tumor management [5, 8, 9].

The coiled-coil-helix-coiled-coil-helix domain-containing protein 2 (CHCHD2), initially described as a regulator of mitochondrial respiration, has recently emerged in the contexts of non-small cell lung and renal cell carcinoma, as well as breast cancer [10-12]. The *CHCHD2* gene is located proximal to *EGFR* on chromosome 7p11.2; as such, *CHCHD2* and *EGFR* are frequently co-amplified in non-small cell lung carcinoma (NSCLC) [10]. *CHCHD2* encodes a 16 kDa protein belonging to a family of 9 evolutionarily conserved small mitochondrial proteins, all containing at least one CHCH domain [13, 14]. The CHCH domain, characterized by two CX9C motifs (two cysteines separated by 9 amino acids), is necessary for simultaneous oxidative folding and protein import into the mitochondrial intermembrane space (IMS) via the CHCHD4-mediated disulfide relay system [15]. CHCHD2 canonically functions as a mitochondrial protein mediating cellular respiration [15-17]. In addition, CHCHD2 has been demonstrated across a variety of biological contexts to regulate other cellular functions, including cell migration and regulation of apoptosis [10, 18, 19]. Furthermore, previous evidence implicates CHCHD2 as a protein involved in mitonuclear communication with the ability to act as a nuclear transcription factor in response to hypoxia, inducing expression of complex IV subunit 4 isoform 2 (*COX4I2*) and itself maximally at 4% O2 [20]. However, the subcellular localization, distribution, and dynamics of CHCHD2 in GBM cells in response to hypoxia has not been described. Additionally, a mechanism governing its mitochondrial export and subcellular redistribution remains elusive. Furthermore, the functional capabilities of CHCHD2 in the context of GBM remain unexplored.

The objective of this study was to characterize the functional capacity of CHCHD2 in GBM cells expressing EGFRvIII, as well as interrogate the intracellular dynamics of CHCHD2 in response to metabolic stressors in U87 and U87vIII GBM cells. Results indicate that subcellular distribution of CHCHD2 between mitochondria and nuclei is sensitive to expression of EGFRvIII and hypoxia, and that CHCHD2 participates in mediating a number of GBM cell functions representing disease hallmarks, which may inspire therapeutic strategies undermining mitochondrial biology to potentially improve GBM tumor management.

## Materials and Methods

### CHCHD2 gene amplification and mRNA expression analysis

Analysis of *CHCHD2* gene amplification patterns across GBM tumors was performed on The Cancer Genome Atlas (TCGA) Provisional GBM database using cBioPortal (http://www.cbioportal.org) [21, 22]. Analysis was limited to tumor samples with available copy number alteration (CNA) data (n = 577). Analysis of CHCHD2 mRNA expression levels (HG-U133A Array) was compared among GBM tumors using publicly available data via Gliovis (http://www.gliovis.bioinfo.cnio.es/) [23]. Tumors were stratified by variables including tumor grade, tumor vs non-tumor tissue, GBM subtype, glioma CpG island methylator phenotype (G-CIMP) status, *IDH1* mutational status, and *MGMT* mutational status.

### Cell culture

The human U87 parental GBM cell line, as well as U87 GBM cells transduced to stably express the constitutively active EGFRvIII mutant (U87vIII) were generously provided by Dr. Nathan Price (Institute for Systems Biology, Seattle, WA). U87 and U87vIII cells transfected to express the pDsRed2-Mito fluorescent mitochondrial marker were used for a subset of immunofluorescence experiments.

GBM patient-derived cells (PDC) serially passaged as orthotopic patient-derived xenografts in mice were generously provided by Dr. Jann Sarkaria (Mayo Clinic, Rochester, MN). Upon receipt, PDCs were cultured in 3D methacrylated gelatin (GelMA) hydrogels without (7% wt gelatin) or with hyaluronic acid (HA) (6% wt gelatin, 1% wt HA). The panel of PDCs analyzed in the current study displayed disparate EGFR/PTEN status, MGMT methylation state (methylated/unmethylated: M/U), molecular subtype (mesenchymal/classical: M/C), invasive characteristics in mouse orthotopic xenografts (0: lo, 7: hi), and sensitivity to erlotnib (0: not sensitive, 100: sensitive) (**Figure 1**) [24].

**Figure 1.**
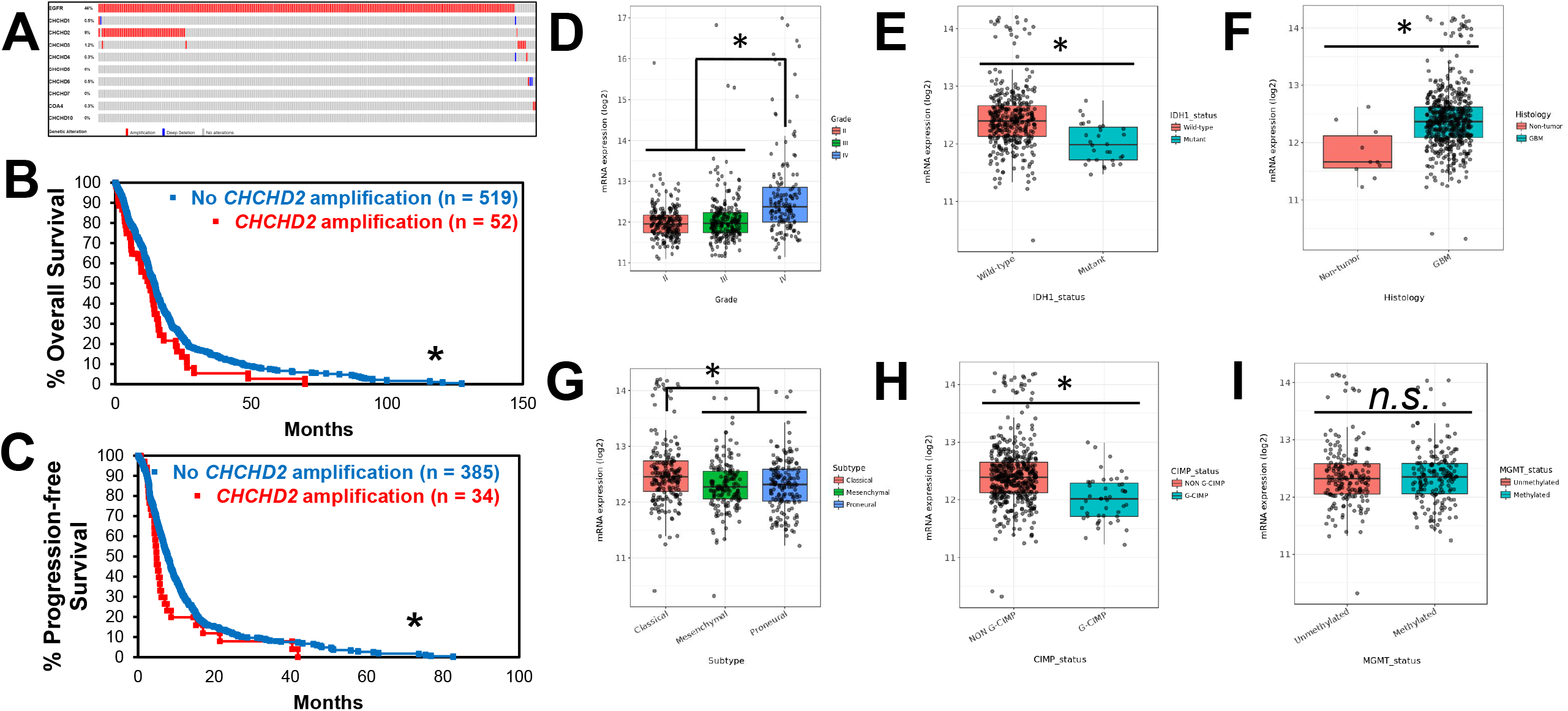
Amplification patterns of CHCHD2 across GBM tumors. **(A)** Oncoprint from cBioPortal (www.cbioportal.org) representing tumors with amplification (red), deep deletion (blue), or no alteration (grey) of query genes. Percentages represent percentage of samples analyzed (tumor samples with copy number alteration data, n = 577) with alteration in given gene. *COA4* encodes CHCHD8. **(B)** Overall survival of patients with (red) and without (blue) *CHCHD2* amplification. **(C)** Progression-free survival of patients with (red) and without (blue) *CHCHD2* amplification. * *p* < 0.05. Using GlioVis, CHCHD2 mRNA expression was determined using Human Genome U133A Array was compared across **(D)** glioma grade, **(E)** *IDH1* mutational status, **(F)** non-tumor vs tumor, **(G)** GBM subtype (classical, mesenchymal, proneural), **(H)** G-CIMP status, which stratifies tumors based on genome-wide DNA methylation status, and **(I)** *MGMT* methylation status. Data were analyzed using Tukey’s Honest Significant Difference or pairwise t-test in GlioVis. * *p* < 0.001.

Cells were cultured in Dulbecco’s modified Eagle’s medium (DMEM) containing 1 mM sodium pyruvate, 15 mM HEPES, non-essential amino acids, 10% fetal bovine serum (FBS), and 1% penicillin/streptomycin. Media devoid of FBS was used for serum deprivation experiments. Media without phenol red was used for immunofluorescence experiments. All cell cultures were tested for mycoplasma contamination prior to orthotopic injection for mouse studies.

### CHCHD2 protein levels in patient-derived cells cultured in GelMA hydrogels

Total protein levels of CHCHD2 were analyzed in a panel of GBM patient-derived cells (PDCs) with disparate EGFR/PTEN status, MGMT methylation state, molecular subtype, invasive characteristics in mouse orthotopic xenografts (0: low, 7: hi), and sensitivity to erlotinib (0: not sensitive, 100: sensitive) (**Figure 1**) [24]. PDCs were cultured in GelMA hydrogels without hyaluronic acid (HA) (7% wt GelMA) or with HA (6% wt GelMA, 1% wt HA).

### Hypoxic cell culture

For hypoxic experiments, medium was pre-equilibrated overnight in a BioSpherix™ hypoxic incubator (BioSpherix, Parish, NY) at designated oxygen concentrations (7%, 4%, and 1%) to account for time required for oxygen-saturated media to equilibrate with the gas atmosphere [25]. Standard culture conditions are designated as normoxia (20% O2). All cells were maintained at 37 °C and 5% CO2.

### Generation of CHCHD2 knockout cells

U87vIII CHCHD2 KO cells were derived using CRISPR-Cas9 genome engineering following published protocols [26]. Suitable target sites within exons of the coding sequence for *CHCHD2* were identified using the online WU-CRISPR design tool [27]. Potential guide RNA (gRNA) oligonucleotides were obtained from Integrated DNA Technologies (IDT, Coralville, IA). Each gRNA sequence was 20 nucleotides in length and directly upstream of the protospacer adjacent motif (PAM) 5’-NGG-3’. Three separate gRNA expression constructs were generated by cloning phosphorylated and annealed gRNA oligos into the BbsI (NEB, Ipswich, MA) site of the pSpCas9(BB)-2A-Puro expression vector (Addgene, Watertown, MA) for co-expression of each sgRNA with the Cas9 endonuclease. The integrity of constructs was confirmed by plasmid sequencing (University of Illinois Urbana-Champaign Roy J. Carver Biotechnology Center).

U87vIII cells were transfected with Cas9-CHCHD2gRNA expression constructs using Lipofectamine 2000 Reagent according to the manufacturer’s protocol (ThermoFisher, Waltham, MA). Stably transfected cells were selected with puromycin (10 µg/mL). Successful CHCHD2 protein knockout was confirmed by western blot as described below. Assessment of genomic mutational status was conducted via nested PCR of the region containing the induced double-strand break using two primer sets. The PCR product was cloned into the pCR2.1-TOPO vector (TOPO TA Cloning Kit, ThermoFisher, Waltham, MA) and confirmed by sequencing (UIUC Roy J. Carver Biotechnology Center).

### Measurement of oxygen consumption rate

Mitochondrial respiration of U87vIII CHCHD2 WT and KO cells was compared using the Seahorse XFp Extracellular Flux Analyzer. Cells were seeded at 1 × 10^4^ cells/well five hours prior to conducting the mitochondrial stress test, according to the manufacturer’s protocol (Agilent, Santa Clara, CA). Serial applications of oligomycin (ATP synthase inhibitor, 1 µM), FCCP (protonophore, 0.5 µM), and rotenone and antimycin A (respiratory complex I and III inhibitor, respectively, 0.5 µM) over time enabled calculation of various parameters of mitochondrial respiration in both cell lines.

### Measurement of compartmentalized glutathione redox poise

U87vIII CHCHD2 WT and KO cells were transfected with genetically encoded, fluorescent redox biosensors targeted to the cytosol (cyto-Grx1-roGFP2) or mitochondrial matrix (mito-Grx1-roGFP2), previously described by our lab [28]. Cells expressing cytosolic or mitochondrial Grx1-roGFP2 were seeded at equal densities in standard culture medium without phenol red in µ-Slide eight-well ibiTreat microscopy chambers (Ibidi, Munich, Germany). Time-lapse images were collected with a fluorescence-enabled inverted microscope (Axiovert 200 M, Carl Zeiss, Feldbach, Switzerland). Dual-excitation imaging of live cells used 395 and 494 nm excitation cubes, and an emission filter at 527 nm was used for both cubes. Exposure times were set to 100-200 ms, and images were taken every 15 s. To assess the effect of GSH synthesis inhibition on GSH:GSSG status, cells were pre-treated with buthionine sulfoximine (BSO, 100 µM) for 24, 48, and 72 h before time-lapse image acquisition. Acquired images were processed with Zeiss Axiovision SE64 Rel6.8 software, via manual selection of three to five individual cells to obtain multiple regions of interest in each time lapse. The means of emission intensities at 527 nm were exported to Excel files and corrected by background subtraction.

### Cell proliferation

Proliferation of U87vIII CHCHD2 WT and KO cells was compared over 72 h in normoxic culture conditions (20% O2) and hypoxia (1% O2) using the sulforhodamine B assay according to published protocols [29]. Briefly, cells were seeded at equal densities in a 96-well plate, followed by fixation with 10% trichloroacetic acid for 1 h at 4° C at 0 h and 72 h timepoints. After washing 4x with water and air-drying at RT, 0.057% sulforhodamine B (SRB) solution (wt/vol in 1% acetic acid) was applied to each well and incubated for 30 min at RT, followed by washing 4x in 1% acetic acid. After drying, bound SRB was solubilized in 10 mM Tris base solution (pH 10.5), and plates were shaken for 30 min at RT. The optical density (OD) of each well was measured at 510 using a BioTek Synergy™ HT microplate reader (BioTek, Winooski, VT). OD values at 72 h were corrected by 0 h OD subtraction to account for possible variations in initial seeding densities.

### Cell cytotoxicity assays

Sensitivity of U87vIII CHCHD2 WT and KO cells to a panel of cytotoxic agents was determined using the sulforhodamine B assay according to published protocols [29]. The cytotoxicity of sulfasalazine (SSZ, xCT cystine-glutamate antiporter inhibitor), erlotinib (Erl, EGFR receptor tyrosine kinase inhibitor), temozolomide (TMZ, DNA alkylating agent), and Pac-1 (procaspase-3 activator) [30] were assessed using dosages derived from the literature.

### Hydrogel preparation and measurement of cell invasion

The invasive behavior of U87vIII CHCHD2 WT and KO cells was examined in normoxic culture conditions (20% O2) and hypoxia (1% O2). Invasion was quantified within methacrylamide-gelatin (GelMA) hydrogels via a bead invasion assay described previously [31-33]. Hydrogels used in this study were made using 5% wt GelMA, ∼53% degree of methacrylamide functionalization determined via H^1^-NMR (data not shown), and photopolymerized under UV light (AccuCure LED 365 nm, 7.1 mW cm^-2^ for 30 s) in the presence of a lithium acylphophinate photoinitiator. The compressive modulus of 5% wt GelMA hydrogels was measured using a Instron 5943 mechanical tester with the Young’s modulus obtained from the linear region of the stress-strain curve (0-10% strain) [33].

To examine invasive behavior, U87vIII CHCHD2 WT and KO cells were seeded onto collagen-coated dextran beads (∼200 µm diameter, GE Life Sciences, Pittsburgh, PA) at a density of 2 × 10^6^ cells per 5 × 10^3^ beads in 5 mL DMEM. Cell-bead suspensions were lightly shaken for one min, every 30 min, for 5 h to facilitate cell adhesion to beads. Cell-coated beads were then encapsulated in pre-polymerized GelMA hydrogel solution, and bead-containing hydrogels were cultured in standard DMEM in either normoxic culture conditions (20% O2) or hypoxia (1% O2) for 7 days. Cell invasion distance was measured from the bead surface using ImageJ from images acquired via fluorescent microscopy after fixing and staining cells with DAPI (Invitrogen, Carlsbad, CA) (10 µg/mL in 1X PBS). Cell invasion is reported as the mean invasion of all cells from the surface of the bead [31].

### Western blot

Total protein levels of CHCHD2 (1:500, NBP1-94106, Novus Biologicals, Centennial, CO)), the glutamate-cystine antiporter xCT (1:500, ab37185, Abcam, Cambridge, UK), GPx-1/2 (1:100, sc-133160, Santa Cruz Biotechnology, Dallas, TX), GPx-4 (1:100, NBP2-75511, Novus Biologicals, Centennial, CO), and matrix metalloproteinase 2 (MMP-2) (1:500, 10373-2-AP, Proteintech Group, Rosemont, IL) were analyzed using western blot. β-actin (1:1000, #4967, Cell Signaling Technology, Danvers, MA) was used as loading control. Total protein concentrations were determined using the Pierce BCA assay (Thermo Fisher, Waltham, MA). Protein lysates were mixed 1:1 with 2X Laemmli Sample Buffer (Bio-Rad, Hercules, CA) (5% β-mercaptoethanol) and heated at 95 °C for 5-10 min. Denatured lysates were loaded into 4-20% Mini-Protean ® TGX™ electrophoresis gels (Bio-Rad, Hercules, CA), and SDS-PAGE was run at 150 V for 1-1.5 h. Proteins were transferred to a nitrocellulose membrane (Amersham, Little Chalfont, UK) at 300 mA for 2 h at 4° C. Membranes were then blocked with either 5% BSA or 5% non-fat dry milk for 1 h at RT, then incubated in primary antibody at designated concentrations overnight at 4° C. Membranes were washed in TBS-T for 5 min 3x, then incubated in HRP-linked goat anti-rabbit secondary antibody (1:2500, #7074, Cell Signaling Technology, Danvers, MA) at RT for 1.5 h. Following TBS-T washes (3x 5 min each), membranes were imaged using the SuperSignal™ West Femto Maximum Sensitivity Substrate (Thermo Fisher, Waltham, MA) in an ImageQuant LAS 4010 (GE Life Sciences, Pittsburgh, PA). Analysis of bands was conducted using ImageJ.

### Animals and orthotopic injection

NOD.Cg-*Prkdc*^*scid*^ *Il2rg*^*tm1Wjl*^/SzJ mice (NOD *scid* gamma (NSG™)) mice (The Jackson Laboratory, Bar Harbor, ME) aged 8-11 weeks were used for this study. Prior to GBM cell induction, mice were anesthetized through intraperitoneal administration of 100 mg/kg ketamine and 10 mg/kg xylazine. The sexes were distributed into groups evenly, followed by injection with U87vIII WT or U87vIII CHCHD2 KO cells. The injection concentration was 1 × 10^5^/μL, with desired volume being 0.5 μL per mouse. A 0.5 μL Hamilton syringe inserted according to the following coordinates in relation to bregma: rostral 0.5 mm, lateral to right 2.25 mm, and 3.3 mm lowered into brain tissue. GBM cells were infused for 30-60 seconds to ensure limited injection backflow, followed by a one minute waiting period until removal of needle. The incision was then closed with a small amount of VetBond. Post-induction the mice were weighed daily to track percent weight change and were additionally scored for neurological symptoms of tumor formation. A weight loss of 20%, severe doming of the skull, rotational spinning, and paralysis were the primary indicators for the mouse to be sacrificed. Post sacrifice, the brain was fixed in 4% paraformaldehyde for 24 hours and sectioned coronally for tumor histology.

### Statistical analysis

Differences among means were tested for using Student’s t-test or one-way ANOVA, followed by post-hoc Student-Newman-Keuls analysis where appropriate. Statistical significance was set at p < 0.05. Variance is reported as standard error of the mean. Odds ratios for *CHCHD2* co-amplification with *EGFR* and log-rank tests for overall and progression-free survival were conducted in cBioPortal [21, 22]. Tukey’s Honest Sigificant Difference or pairwise t-tests were conducted to compare mRNA expression levels in GlioVis [23].

## Results

CHCHD2 *is amplified in a subset of GBM tumors and is associated with decreased patient survival* Analysis of *CHCHD2* amplification patterns across GBM tumors was performed on The Cancer Genome Atlas (TCGA) Provisional GBM database using cBioPortal (http://www.cbioportal.org) [21, 22]. Analysis was performed on tumor samples with available copy number alteration (CNA) data (n = 577). Amplification of *CHCHD2* was observed in 9% of GBM tumors (**Figure 1A**). Of tumors with *EGFR* amplification, *CHCHD2* was co-amplified in 20% of cases, with a significant tendency to co-occur (log2 odds ratio > 3, p < 0.001). Of the 9 proteins in the CHCH domain-containing protein family, only *CHCHD2* was amplified at an appreciable frequency in GBM tumors, with the next most frequently amplified CHCH protein being *CHCHD3* (1.2%) (**Figure 1A**). Additionally, patients with *CHCHD2*-amplified tumors displayed decreased overall survival (OS, 12.48 mo vs 14.45 mo, log-rank p = 0.0223) (**Figure 1B**) and progression-free survival (PFS, 4.86 mo vs 7.82 mo, log-rank p = 0.0420) (**Figure 1C**). While this effect could potentially be explained by the propensity for *EGFR* to co-amplify with *CHCHD2*, amplification of *EGFR* was not solely associated with decreased patient survival (**Supplementary Fig. S1**). These data indicate that amplification of the *CHCHD2* gene occurs in a subset of GBM tumors, is associated with decreased OS and PFS, and is nearly always accompanied by *EGFR* amplification.

Using the GlioVis data portal, CHCHD2 mRNA expression was observed to be increased in grade IV gliomas (GBM) relative to grade II and III (**Figure 1D**), as well as in *IHD1*-wt tumors (primary GBM) compared to *IDH1*-mutant (secondary GBM) (**Figure 1E**). CHCHD2 expression was also increased in GBM tumor compared to non-tumor tissue (**Figure 1F**). Additionally, CHCHD2 expression was increased in classical subtype tumors (**Figure 1G**), likely due to the location of *CHCHD2* on chromosome 7, the amplification of which defines classical GBM tumors and is accompanied by a focused predilection for *EGFR* amplification [34]. Tumors exhibiting genome-wide promoter hypermethylation, i.e. glioma-CpG island methylator phenotype (G-CIMP), displayed decreased CHCHD2 mRNA expression (**Figure 1H**), consistent with epigenetic methylation-induced silencing of *CHCHD2*. No differences in CHCHD2 expression were observed between *MGMT* methylated vs non-methylated tumors (**Figure 1I**).

### CHCHD2 protein levels vary across GBM patient-derived cell samples

Total protein levels of CHCHD2 were analyzed in a panel of GBM patient-derived cells (PDCs) cultured in GelMA hydrogels (7% wt gelatin). PDCs were characterized by disparate EGFR/PTEN status, MGMT methylation state, molecular subtype, invasive characteristics in mouse orthotopic xenografts (0: low, 7: hi), and sensitivity to erlotinib (0: not sensitive, 100: sensitive) (**Figure 2A**). Included in this panel were: GBM10 (EGFR^-^/PTEN^-^), GBM44 (EGFR^-^/PTEN^-^), GBM12 (EGFR^+^/PTEN^wt^), GBM39 (EGFRvIII/PTEN^wt^), and GBM6 (EGFRvIII/PTEN^wt^). Notably, of PDCs analyzed in this study, CHCHD2 levels were greatest in GBM6, which expresses EGFRvIII, is the most invasive, is relatively resistant to erlotinib, and is of the classical subtype (**Figure 2B**). These results are congruent with mRNA levels of CHCHD2 being greatest in classical subtype tumors (**Figure 1G**).

**Figure 2.**
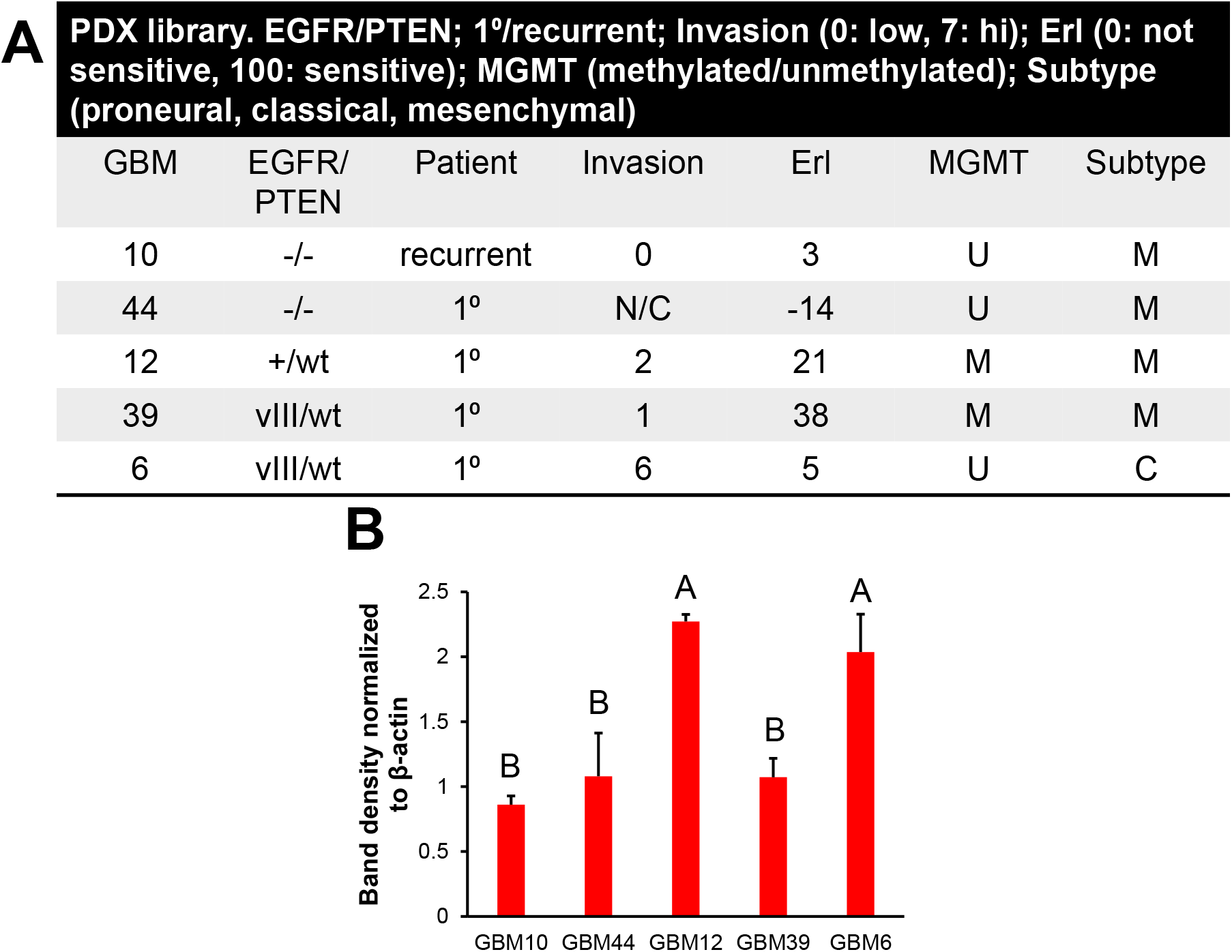
CHCHD2 protein levels in GBM patient-derived cells. **(A)** Patient-derived xenograft (PDX) subset analyzed in the current study, with descriptors including EGFR/PTEN status, patient status, invasion in mouse orthotopic xenografts, erlotinib sensitivity, MGMT methylation status, and subtype. **(B)** CHCHD2 total protein levels in PDC samples from each PDX cultured in GelMA hydrogels containing matrix-immobilized HA (6% wt GelMA, 1% wt HA). Differing superscript: *p* < 0.05. * denotes significant effect of HA, *p* < 0.05

### Knockout of CHCHD2 alters mitochondrial respiration in U87vIII cells

The gene amplification, mRNA, and protein expression patterns of CHCHD2 observed across clinically relevant GBM patient samples suggested a biologically relevant role for CHCHD2 in mediating GBM cell phenotypes. To dissect the essential functionality of CHCHD2 in the context of GBM, U87vIII CHCHD2 KO cells were derived using CRISPR-Cas9 following published protocols (**Supplementary Fig. S2**) [26, 27], and protein knockout validated at the protein level using western blot (**Figure 3A**). CHCHD2 was first described during a computational screen as a mediator of mitochondrial respiration [16]. To determine if CHCHD2 KO impacted mitochondrial respiration in GBM cells, a mitochondrial stress test was conducted on U87vIII CHCHD2 WT and KO cells using a Seahorse XFp Extracellular Flux Analyzer (**Figure 3B**) [35]. CHCHD2 KO cells displayed decreased basal and maximal oxygen consumption rate (OCR) (**Figure 3B-D**). Spare respiratory capacity was also significantly lower in CHCHD2 KO cells (**Figure 3E**), indicating a decreased ability of U87vIII CHCHD2 KO cells to respond to increased energy demand. Additionally, the amount of oxygen consumed coupled to ATP production by ATP synthase was significantly lower in CHCHD2 KO cells (**Figure 3F**). Collectively, these data indicate that CHCHD2 is indeed required for efficient mitochondrial respiration in U87 GBM cells expressing EGFRvIII.

**Figure 3.**
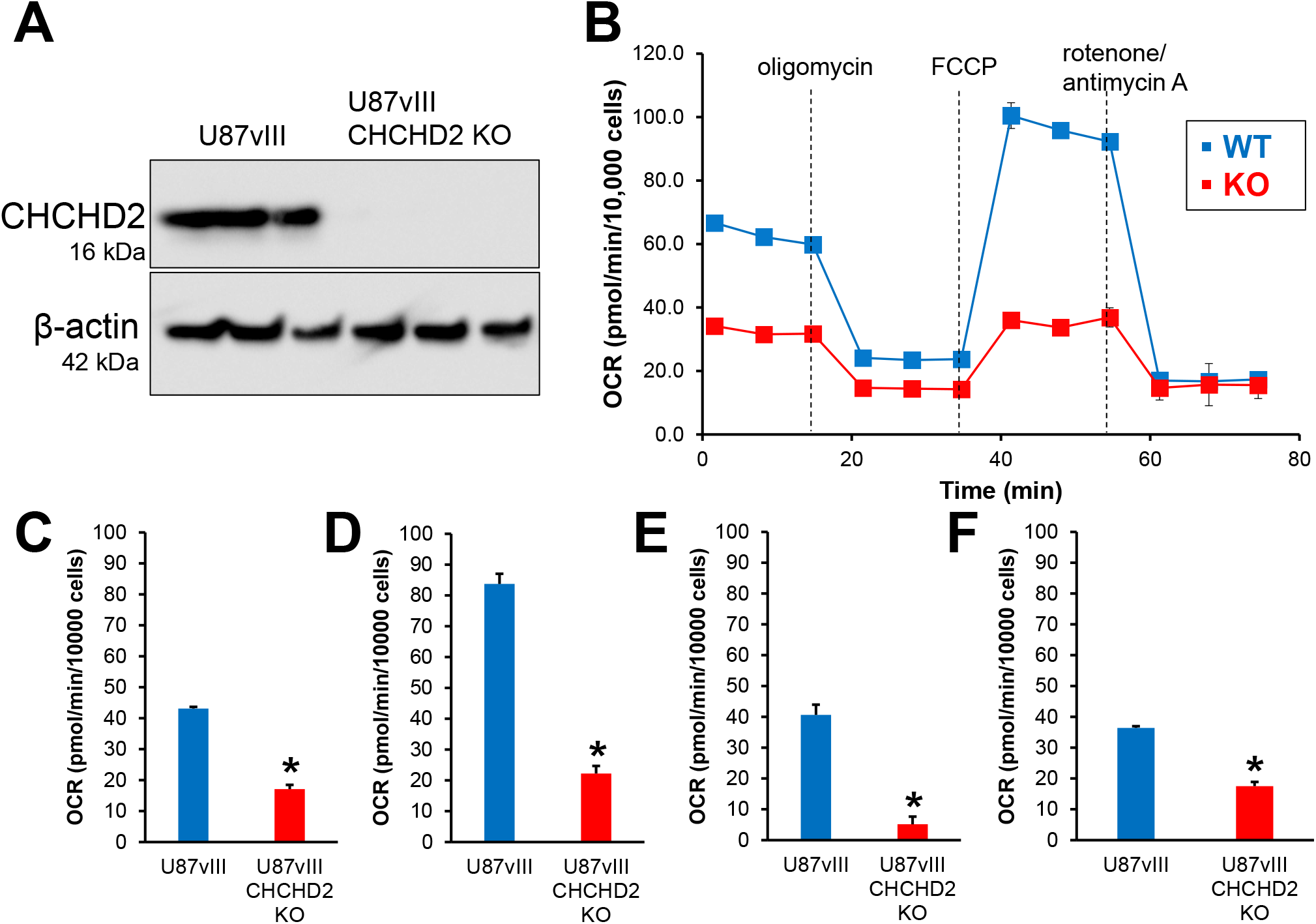
Mitochondrial respiration in U87vIII and U87vIII CHCHD2 KO cells. **(A)** Western blot of U87vIII and U87vIII CHCHD2 KO cells. **(B)** OCR traces of U87vIII CHCHD2 WT and KO cells during mitochondrial stress test. **(C)** Basal OCR, **(D)** maximal respiration, **(E)** spare respiratory capacity, and **(F)** OCR coupled to ATP production in U87vIII CHCHD2 WT and KO cells. Data are presented as mean ± SE. * *p* < 0.05.

### Knockout of CHCHD2 leads to a more reduced glutathione redox pool in the mitochondrial matrix

Upon observing defects in mitochondrial respiration, it was further hypothesized that deficient electron transport chain function would result in a more oxidized intracellular redox environment manifested by increased amounts of oxidized glutathione (GSSG). Glutathione (GSH), an enzymatically produced tripeptide of cysteine, glycine, and glutamate, is the main intracellular redox buffer, which, along with thioredoxins and glutaredoxins, maintains thiol redox status [36]. The glutathione redox couple (reduced and oxidized glutathione, GSH and GSSG respectively), along with glutathione peroxidase (GPx) and glutathione reductase (GR), comprises the glutathione system, which maintains thiol redox homeostasis and functions in antioxidant defense [37]. The balance of reduced to oxidized glutathione (GSH:GSSG) thus represents intracellular redox status, and can be interrogated within live cells using genetically encoded, fluorescent redox biosensors (Grx1-roGFP2) (**Figure 4A**). Additionally, such probes can be targeted to various subcellular compartments, including cytosol (cyto-Grx1-roGFP2) or mitochondrial matrix (mito-Grx1-roGFP2) to measure compartmentalized GSH redox status in live cells in real time via ratiometric fluorescence intensity measurements (**Figure 4B**) [28]. The percent of oxidized mito-Grx1-roGFP2 in U87vIII CHCHD2 KO cells was decreased by 15.8% compared to that measured in CHCHD2 WT cells, indicating that the pool of glutathione in the mitochondrial matrix of KO cells is more reduced than WT counterparts (**Figure 4C**). This effect was confined to the mitochondrial matrix, as the cytosolic glutathione pool was not affected by CHCHD2 KO (**Figure 4D**). Treatment with buthionine sulfoximine (BSO), an inhibitor of glutamate-cysteine ligase (GCL, the rate-limiting enzyme in GSH synthesis), led to similar oxidation of the glutathione pool over time in the mitochondrial matrix of both U87vIII CHCHD2 WT and KO cells (**Figure 4E**). Additionally, levels of components of the glutathione system, including the glutamate-cystine antiporter xCT, GPx-1/2, and GPx-4, were all unaltered in CHCHD2 KO cells (**Figure 4F**). These data demonstrate a role for CHCHD2 in mediating glutathione redox balance specifically in the mitochondrial matrix, albeit through a mechanism not involving GSH synthesis, cystine import through xCT, or reduced GSH flux through glutathione peroxidases 1, 2, or 4.

**Figure 4.**
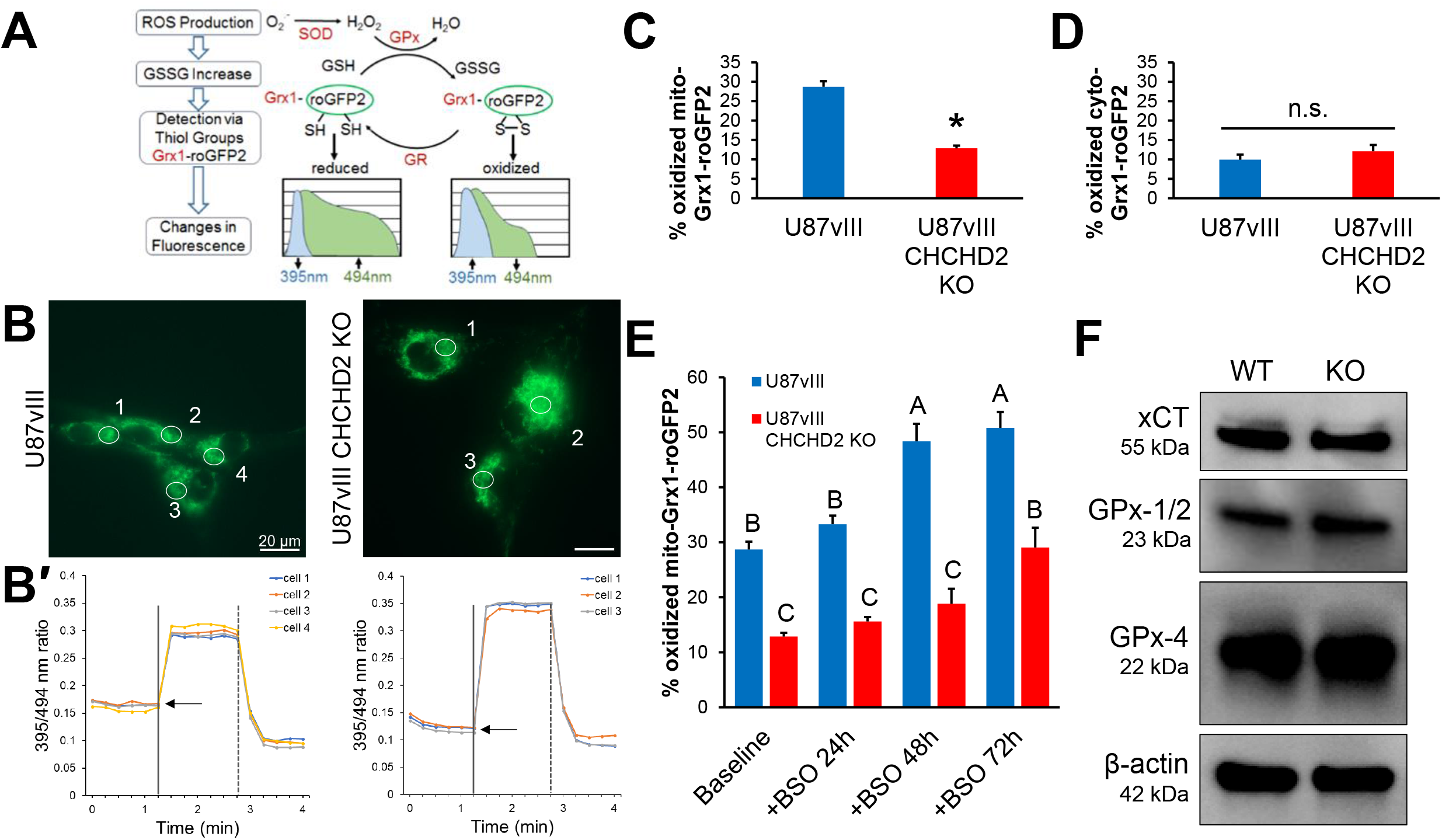
Glutathione redox poise in U87vIII CHCHD2 WT and KO cells. **(A)** Schematic of the molecular mechanism of the Grx1-roGFP2 sensor and redox response of the compartmentalized probe to exogenous oxidant and reductant. Superoxide (^•^O2−) is rapidly converted by superoxide dismutase (SOD) into H2O2, which is then reduced by glutathione peroxidase (GPx) to water. Glutaredoxin (Grx) fused to roGFP2 efficiently and rapidly equilibrates the probe with alterations in the local GSH:GSSG ratio. Additionally, thiol-disulfide equilibration is reversible, as GSH reductase (GR) catalyzes reduction of GSSG to GSH. **(B)** Representative fluorescence images demonstrate the sensor targeted to mitochondria of U87vIII CHCHD2 WT (left) and KO cells (right). **B’**. Corresponding time-lapse responses of the 395/494 nm ratio to treatment with 1 mM diamide (vertical solid line) to the fully oxidized state and 10 mM DTT (vertical dashed line) to the fully reduced state. Each trace designates a separate cell. Arrows represent basal oxidation level of probe. **(C)** Percent oxidized mito-Grx1-roGFP2 and **(D)** cyto-Grx1-roGFP2 in U87vIII CHCHD2 WT and KO cells. **(E)** Percent oxidized mito-Grx1-roGFP2 in U87vIII CHCHD2 WT and KO cells at baseline, and after treatment with GSH synthesis inhibitor BSO for 24, 48, and 72 h. **(F)** Western blot of xCT, GPx-1/2, and GPx-4 in U87vIII CHCHD2 WT and KO cells. Data are presented as mean ± SE. *, differing superscript: *p* < 0.05.

### U87vIII cell growth and invasion are negatively impacted by CHCHD2 KO in both normoxia and hypoxia

The observed deficiencies in mitochondrial respiration led to the hypothesis that U87vIII GBM cell growth would be negatively impacted by CHCHD2 KO. To test this hypothesis, U87vIII CHCHD2 WT and KO cells were incubated in either standard oxygen culture conditions (normoxia, 20% O2) or pathophysiologically relevant hypoxia (1% O2). Utilizing the sulforhodamine B (SRB) colorimetric assay [29] to assess cell growth over 72 h, CHCHD2 WT cell growth was observed to be significantly increased in hypoxia (**Figure 5A**). CHCHD2 KO cell growth was decreased compared to normoxic control (**Figure 5A**). Additionally, the growth-inducing effect of hypoxia on U87vIII cells was abrogated upon CHCHD2 KO (**Figure 5A**). These results demonstrate that CHCHD2 is involved in not only mediating U87vIII cell growth in normoxia, but also plays a role in the initial increased cell proliferation observed in hypoxic cells.

**Figure 5.**
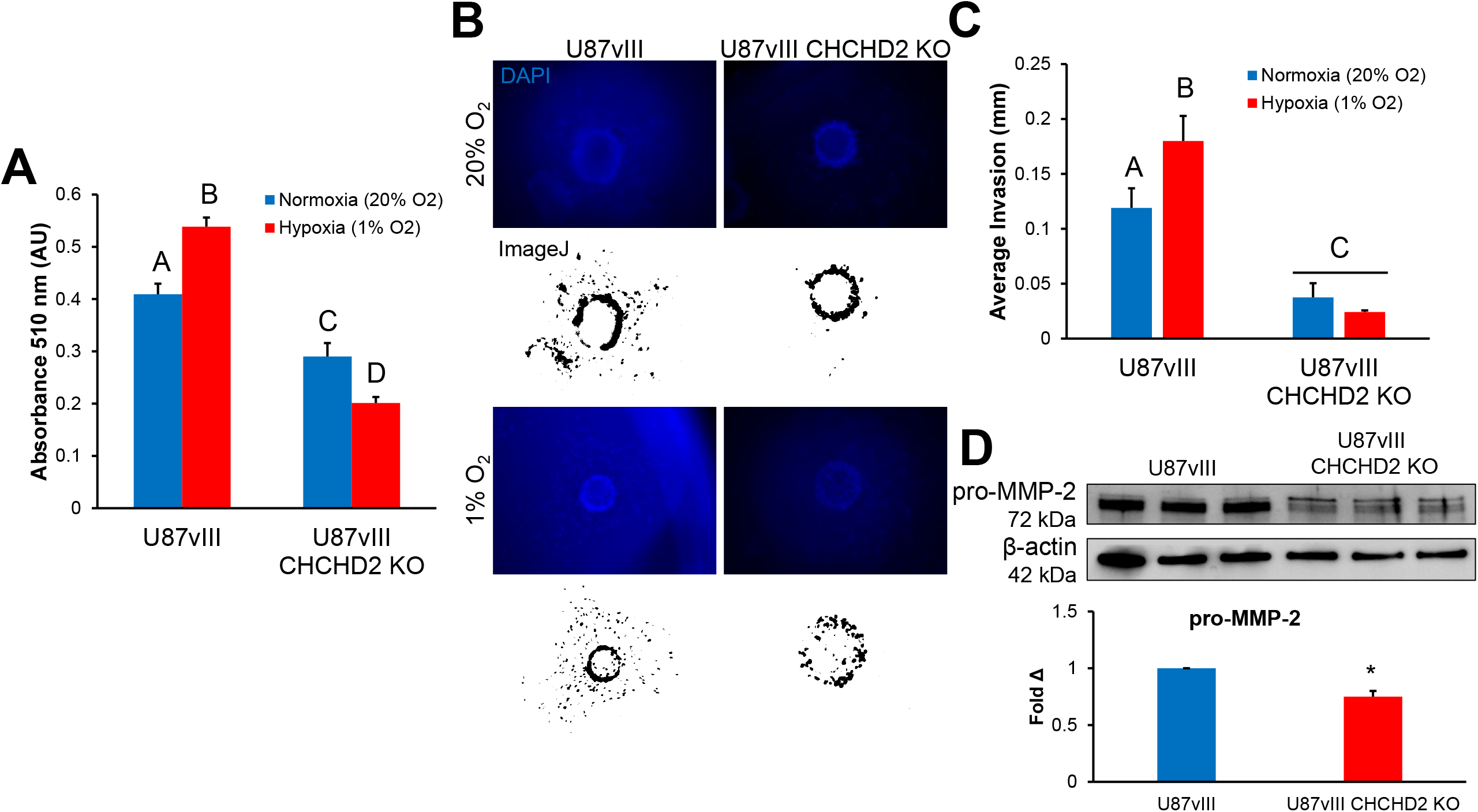
Growth and invasion of U87vIII and U87vIII CHCHD2^KO^ cells in hypoxia. **(A)** Cell growth of U87vIII and U87vIII CHCHD2KO cells over 72 h in standard oxygen conditions and hypoxia determined using the SRB assay. **(B)** Representative images of U87vIII and U87vIII CHCHD2^KO^ cells stained for DAPI after incubation in 21% or 1% O2 for 7 days during bead invasion assay. Associated ImageJ-derived images for average invasion distance analysis are displayed below corresponding fluorescence images. **(C)** Quantification of U87vIII and U87vIII CHCHD2^KO^ cell invasion as determined by the bead invasion assay. **(D)** Western blot of pro-MMP-2 in CHCHD2 WT and KO cells and fold-change normalized to β-actin loading control. Data are presented as mean ± SE. Differing superscript, *: *p* < 0.05.

Rapid, diffuse invasion of GBM cells into tumor margins and into surrounding brain parenchyma represents a major obstacle impeding effective tumor treatment. To determine whether CHCHD2 knockout impacted GBM cell invasion, a bead invasion assay in methacrylated gelatin (GelMA) hydrogels (Young’s modulus 2.9 ± 0.45 kPa) was used as a three-dimensional culture platform to model biophysical aspects of the brain parenchyma and GBM cell invasion [31, 32]. This technique enables monitoring of cell invasion from a defined starting point in a spatiotemporal manner in normal as well as hypoxic three-dimensional culture conditions (**Figure 5B**). Hypoxia (1% O2) stimulated U87vIII cell invasion over long-term culture (7 d) (**Figure 5C**). In contrast, U87vIII CHCHD2 KO cells displayed minimal cell invasion in both normoxia and hypoxia, with the hypoxia-induced invasion observed in CHCHD2 WT cells abrogated (**Figure 5C**). Furthermore, CHCHD2 KO cells displayed decreased basal levels of pro-MMP-2 (matrix metalloproteinase 2) (**Figure 5D**), a key protein involved in the breakdown of extracellular matrix. These data demonstrate a role for CHCHD2 in mediating U87vIII cell invasion, particularly in response to hypoxia.

### CHCHD2 knockout increases U87vIII sensitivity to a variety of cytotoxic drugs

To determine the effect of CHCHD2 KO on cellular resistance to various drugs, cells were treated with increasing concentrations of a panel of agents, and cytotoxicity was assessed using the SRB assay [29]. Included in this panel were: temozolomide (TMZ), a DNA-alkylating agent and the standard-of-care chemotherapy administered to patients with GBM [38]; erlotinib (Erl), a receptor tyrosine kinase inhibitor that inhibits EGFR and EGFRvIII tyrosine kinase activity [8]; sulfasalazine (SSZ), an inhibitor of the cell membrane xCT antiporter, which couples the export of the amino acid glutamate with the import of cystine, thus depriving cells of the rate-limiting substrate to synthesize reduced GSH [39, 40]; and Pac-1, a novel activator of apoptosis which acts on procaspase-3 [30]. Results, shown in **Figure 6**, demonstrate that CHCHD2 KO cells were more susceptible to treatment with TMZ, Erl, and most significantly, SSZ. However, CHCHD2 KO had no effect on cellular sensitivity to treatment with Pac-1 (**Figure 6D**), consistent with previous work demonstrating that CHCHD2 regulates apoptosis upstream of procaspase-3 activity in the apoptotic cascade [19]. These results demonstrate a role for CHCHD2 in mediating cell sensitivity to various drugs relevant to GBM treatment, highlighting CHCHD2 as a promising avenue for future investigation.

**Figure 6.**
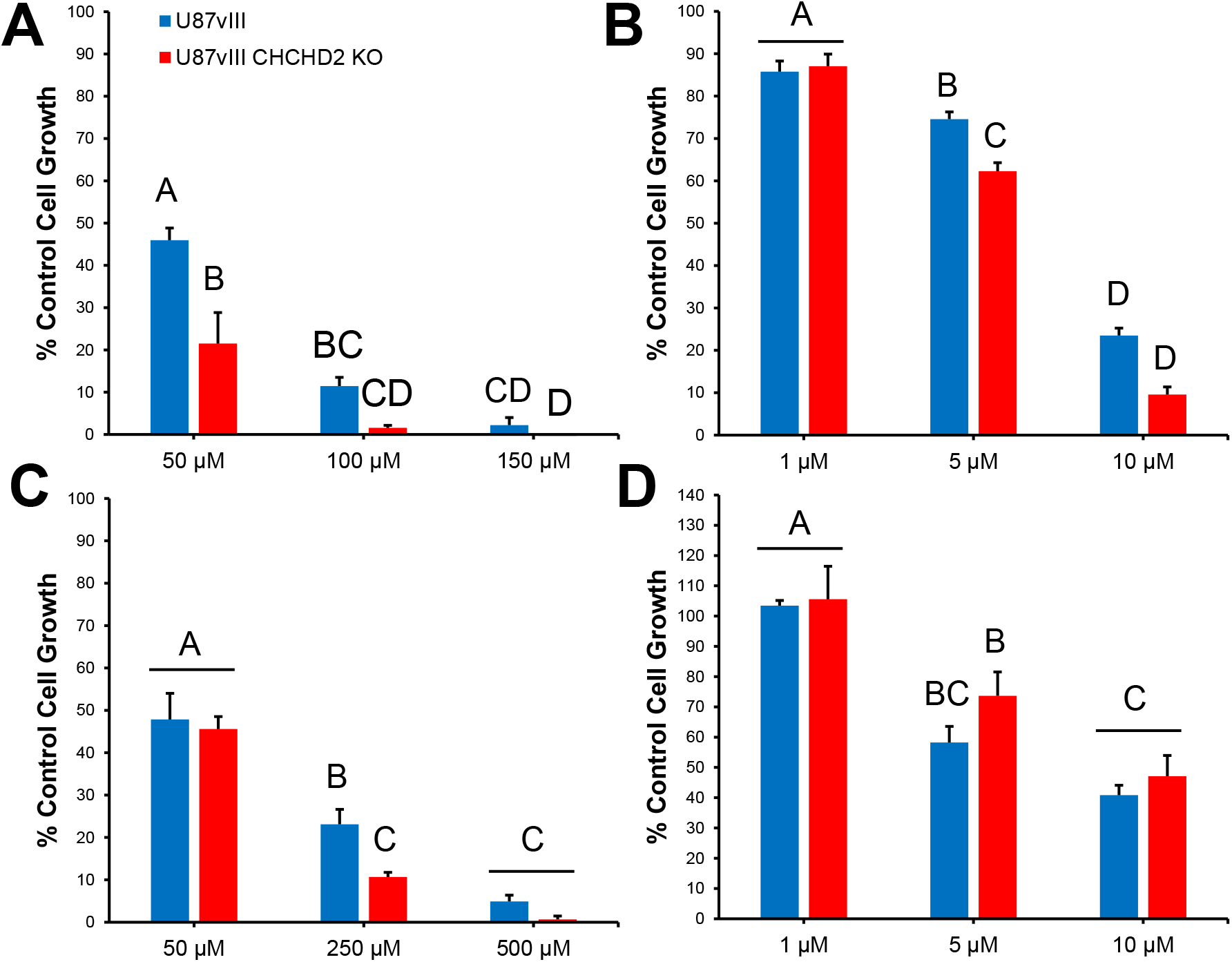
Therapeutic sensitivity of U87vIII and U87vIII CHCHD2^KO^ cells. Sensitivity of U87vIII and CHCHD2KO cells to **(A)** sulfasalazine, **(B)** erlotinib, **(C)** temozolomide, and **(D)** Pac-1 determined using the SRB assay. Data are presented as percent of untreated control, mean ± SE. Differing superscript: *p* < 0.05.

### CHCHD2 knockout impacts mouse overall survival

To determine the effect of CHCHD2 on tumor growth in vivo, NSG mice were orthotopically injected with U87vIII WT and CHCHD2 KO cells. Mice bearing U87vIII WT tumors presented with a median survival of 17 days versus 25 days for U87vIII CHCHD2 KO tumor-bearing mice (**Figure 7A**). Representative coronal sections of mice sacrificed on day 16 post-injection demonstrated substantial tumor growth and infiltration into surrounding brain (**Figure 7B**). Mice injected with U87vIII WT cells displayed rapidly diminishing health by day 13 of the study as evidenced by decreasing weight change (**Figure 7C**). Furthermore, U87vIII WT cells consistently produced tumor occupying a larger percentage of parenchyma compared to CHCHD2 KO-derived tumors (**Figure 7D**). Overall, these data indicate a role for CHCHD2 in promoting progression of GBM tumors in mice.

**Figure 7.**
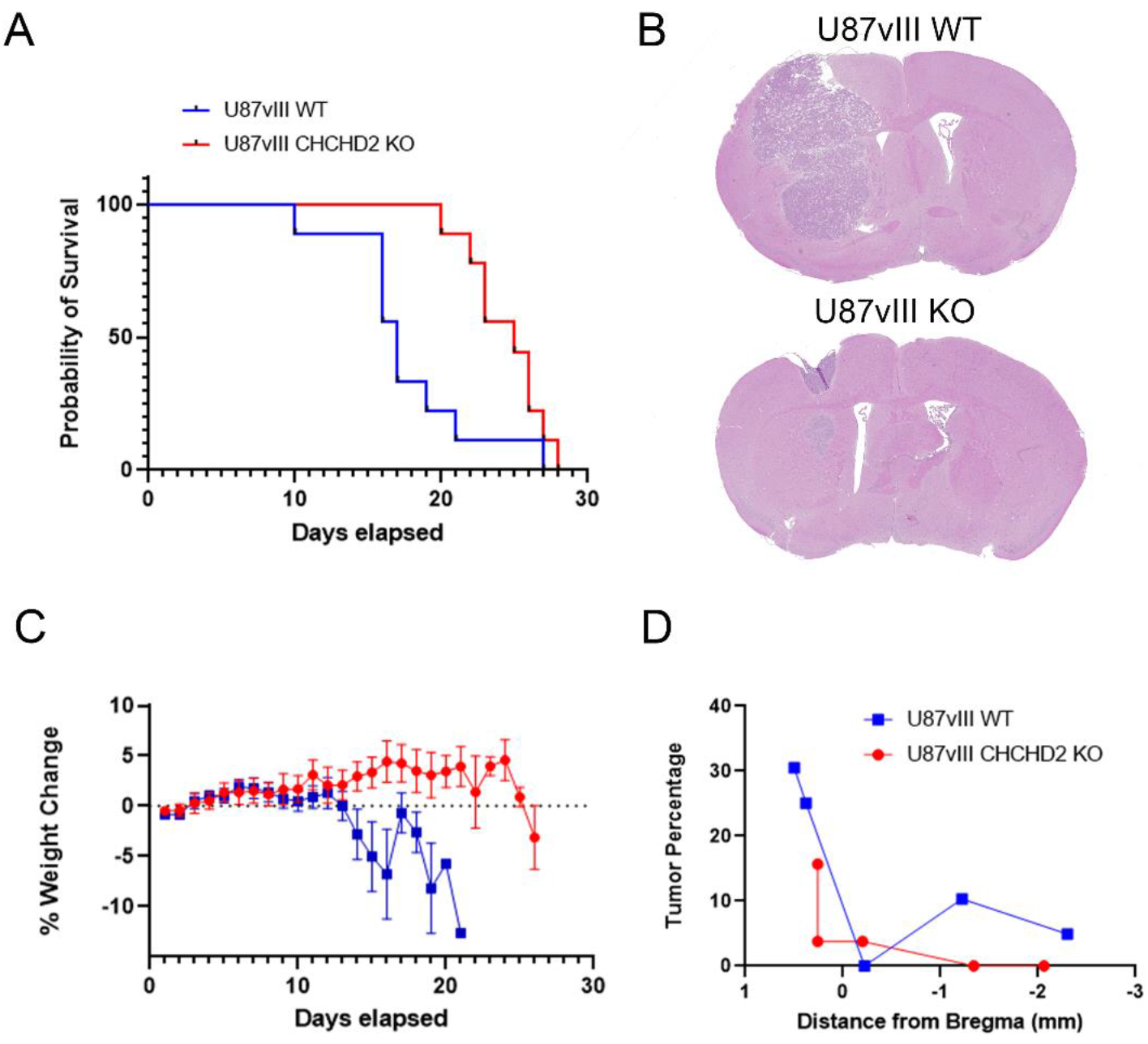
U87vIII CHCHD2 KO-bearing mice displayed greater survival and lower tumor burden relative to mice harboring wild-type U87vIII tumors. **(A)** Survival of tumor-bearing mice injected with U87vIII (blue curve) or U87vIII WT CHCHD2 KO (red curve), with median survival times being 25 days for KO and 17 days for WT mice. **(B)** Representative coronal sections of mice sacrificed on day 16 post-implantation. Sections were cut with respect to bregma: AP = 0.5 mm, LV = 2.25 mm, DV = -3.3 mm. **(C)** Percentage weight change with respect to days elapsed, mean ± SEM. **(D)** Percentage of tumor occupancy in relation to the coronal brain section.

## Discussion

Mitochondria, in addition to regulating cellular energy conservation, also serve as signaling organelles. The vast majority of mitochondrial proteins are encoded by nuclear genes, necessitating the ability for mitochondria to communicate their status to the nucleus in response to metabolic perturbations. Disturbances in ATP and ROS production, damage to mitochondrial DNA, and aberrations in mitochondrial protein folding induce a mitonuclear retrograde signaling pathway by which mitochondria communicate with the nucleus to induce changes in nuclear gene expression in order to maintain metabolic homeostasis [41]. Several proteins have been demonstrated to participate in inter-organelle signaling between mitochondria and the nucleus, including p53, fumarase, the pyruvate dehydrogenase complex, and CHCHD2 [15, 42-44]. CHCHD2 presented as a promising avenue to pursue, given: 1) its proximity to and frequency of co-amplification with *EGFR* with NSCLC and GBM (**Figure 1**) [10], both of which have been characterized as relatively oxidative versus fermentative tumors [45]; 2) its mRNA expression patterns across glioma grade and tumor versus non-tumor tissue (**Figure 1**); 3) its reported oxygen-sensitive transcription factor activity [15, 20]; and 4) its pleiotropic roles mediating cellular functions reminiscent of cancer hallmarks, including proliferation, migration and invasion, and inhibition of apoptosis [15, 18, 19]. Here, we demonstrate that CHCHD2 is involved in mediating therapeutic sensitivity as well as cell growth and invasion in vitro and in vivo in GBM cells expressing EGFRvIII.

Knockout of CHCHD2 in U87vIII GBM cells resulted in decreased baseline respiration as well as spare respiratory capacity (**Figure 3**), an effect corroborated by multiple studies which have characterized CHCHD2 as a canonical regulator of mitochondrial respiration [10, 15-17]. While metabolic reprogramming towards increased glycolytic flux to favor increased cell proliferation is a recognized hallmark of cancer, functional mitochondria remain essential in maintaining malignant cell bioenergetics in particular tumors, including those of lung and brain [45-47]. Notably, mitochondrial spare respiratory capacity has been positively associated with glioma stem cell resistance to radiotherapy [48]. Thus, the decreased spare respiratory capacity measured in U87vIII CHCHD2 KO cells (**Figure 3**) may partially contribute to their increased sensitivity to treatment with TMZ, Erl, and SSZ (**Figure 6**).

Additionally, CHCHD2 has been demonstrated by others to mediate mitochondrial outer membrane permeabilization (MOMP) [19], the “point of no return” during the intrinsic pathway of apoptosis. Indeed, U87vIII CHCHD2 KO cells exhibited increased sensitivity to TMZ, Erl, and SSZ. The observation that no increased in cytotoxicity was seen in CHCHD2 KO cells treated with Pac-1, an activator of procaspase-3 which acts downstream of MOMP in the apoptotic cascade [30], further corroborates the demonstrated role of CHCHD2 early in apoptosis regulation.

Notably, as evidenced by the use of mito-Grx1-roGFP2, the mitochondrial GSH pool was more reduced in response to CHCHD2 KO compared to WT, an effect independent of GSH biosynthesis (**Figure 4**). It should be noted that roGFP2 itself is not directly oxidized by ROS species, but rather equilibrates with the local GSH redox potential, which is influenced by the GSH:GSSG ratio, and in turn is influenced by activity of various ROS-scavenging enzymes that use GSH as a cofactor. In HEK293 cells, CHCHD2 knockdown was accompanied by decreased expression of superoxide dismutase 2 and loss of glutathione peroxidase expression [15]. However, we did not observe changes in protein levels of GPx-1/2, Gpx-4, nor the glutamate-cystine antiporter xCT (**Figure 4**).

Incubation in hypoxia increased U87vIII CHCHD2 WT cell proliferation over 3 days, and increased invasion over 7 days (**Figure 5**). Using the GelMA hydrogel platforms described here, previous work demonstrated that hypoxic U87vIII cells exhibited increased cell proliferation until day 5, at which point cell proliferation stalled, while invasion continued to increase up to day 7 [33]. Notably, CHCHD2 KO abrogated the increased cell growth and invasion in hypoxia exhibited by WT cells (**Figure 5**). The observed decrease in U87vIII CHCHD2 KO cell growth is not likely due to an increase in apoptosis, as evidence has indicated that shRNA-mediated CHCHD2 knockdown does not alter cellular levels of poly (ADP-ribose) polymerase (PARP) [15]. Deficiencies in mitochondrial respiration and ATP production are likely responsible for hampering cell proliferative capacity, and may also be implicated in the repression of cell invasion, as migration and invasion throughout parenchyma is an energy-expensive process, particularly when moving through dense extracellular matrices [49]. The observed decrease in pro-MMP-2 expression in CHCHD2 KO cells provides further explanation for their decrease in invasive capacity (**Figure 5**). Other work has highlighted CHCHD2 as an activator of NIH3T3 fibroblast migration via the AKT-RhoA/ROCK-JNK cascade[18]. The effects of CHCHD2 in vitro were closely mirrored in vivo as well, as mice orthotopically injected with U87vIII CHCHD2 KO cells presented with improved survival time and decreased tumor burden (**Figure 7**).

This study is the first to describe the functional relevance of CHCHD2 in GBM. These data indicate that CHCHD2 is essential for mitochondrial respiration and maintenance of mitochondrial GSH status in U87vIII cells. Additionally, these studies demonstrate a critical role for CHCHD2 in mediating cell growth and invasion in normoxia and hypoxia and resistance to various cytotoxic agents, underscoring CHCHD2 as a mediator of GBM malignant phenotype. Functional outcomes investigated here relate primarily to mitochondrial functions of CHCHD2. As a protein implicated in mitonuclear communication in response to hypoxia, the nuclear functions of CHCHD2 in GBM cells and the panel of nuclear genes it regulates are of equal importance and areas of future work. The nuclear function of CHCHD2 has been demonstrated by others to act in concert with other transcription factors, namely recombining binding protein suppressor of hairless (RBPJ) [20], the main downstream effector protein of the Notch signaling pathway, which itself has been implicated in the maintenance of glioma stem cell maintenance and viability [50]. Future work should seek to identify the complement of genes regulated by CHCHD2 in both normal oxygen conditions as well as hypoxia to provide a deeper understanding of CHCHD2 function in GBM.

**Supplementary Fig. S1**. EGFR gene amplification and survival in GBM cBioPortal

**Supplementary Fig. S2. CRISPR-Cas9-mediated knockout of CHCHD2 in U87vIII cells. (A)** The workflow for the derivation of CHCHD2 KO cells using CRISPR-Cas9, as well as the figure above, were based on and adapted from published work by Ran et al. Three potential gRNAs were designed using the online sgRNA design tool WU-CRISPR (http://crispr.wustl.edu/), ligated into separate linearized pSpCas9(BB)-2A-puro vectors, and recombinant plasmids transfected into separate U87vIII cell populations. Only sgRNA 2 (orange arrow) successfully yielded U87vIII CHCHD2^KO^ cells. **(B)** Primer sets for nested PCR of CHCHD2 genome modification sequencing. **(C)** Genomic screening for induced mutations at the site of interest targeted by the sgRNA. Cyan: internal primer targets for nested PCR (set 2). Yellow: 20 bp sequence within *CHCHD2* exon 2 targeted by sgRNA. Green: protospacer adjacent motif (PAM), required for Cas9 binding to target site. Red: stop codons accompanying insertions (gray).

## Acknowledgements

We would like to further acknowledge Courtney Tank for assistance with experiments.

