## Supplemental Figures 1-2 for "CHCHD2 mediates glioblastoma cell proliferation, mitochondrial metabolism, hypoxia-induced invasion, and therapeutic resistance"

### Slide 1
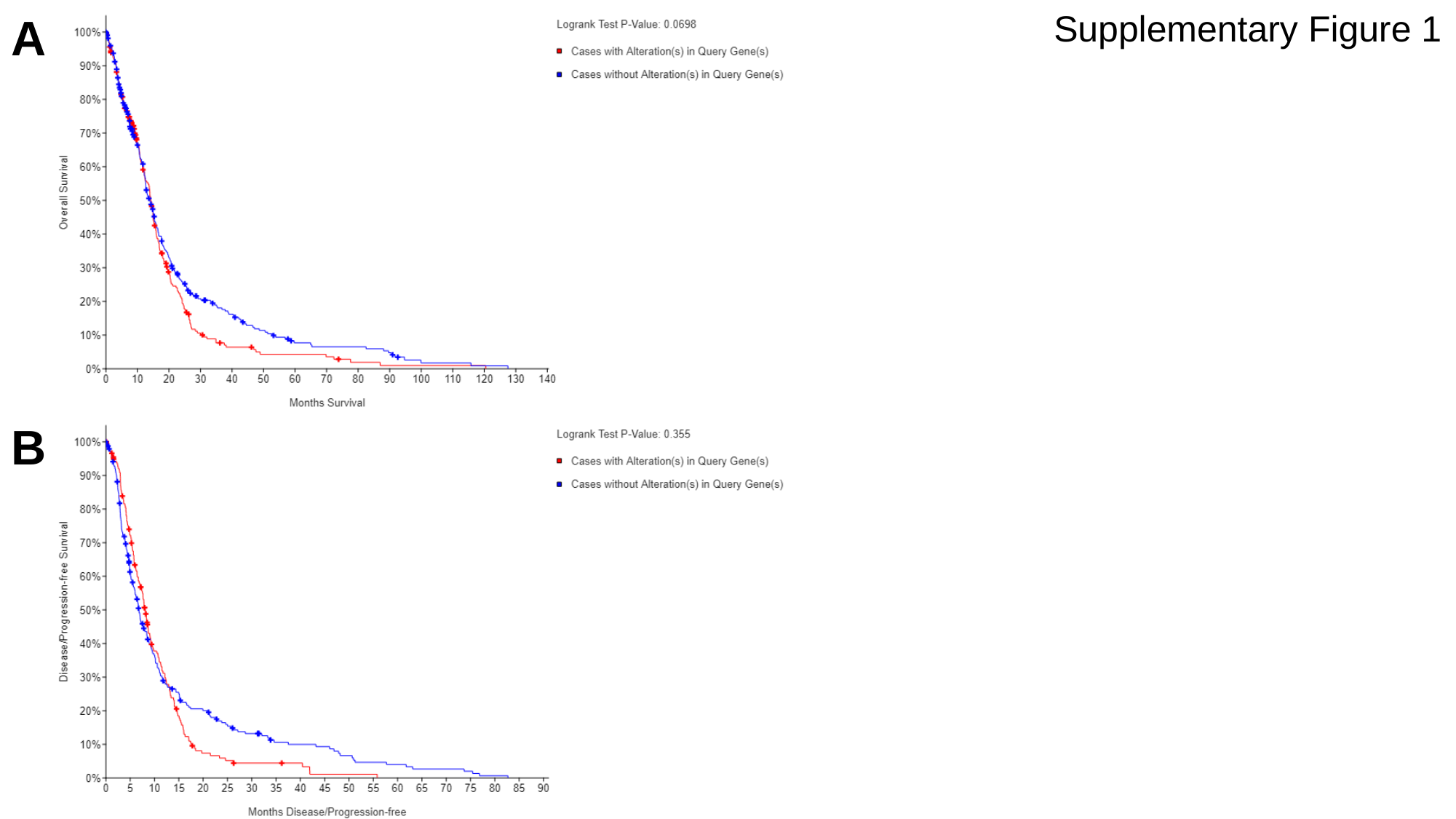

Supplementary Figure 1
A
B

### Slide 2
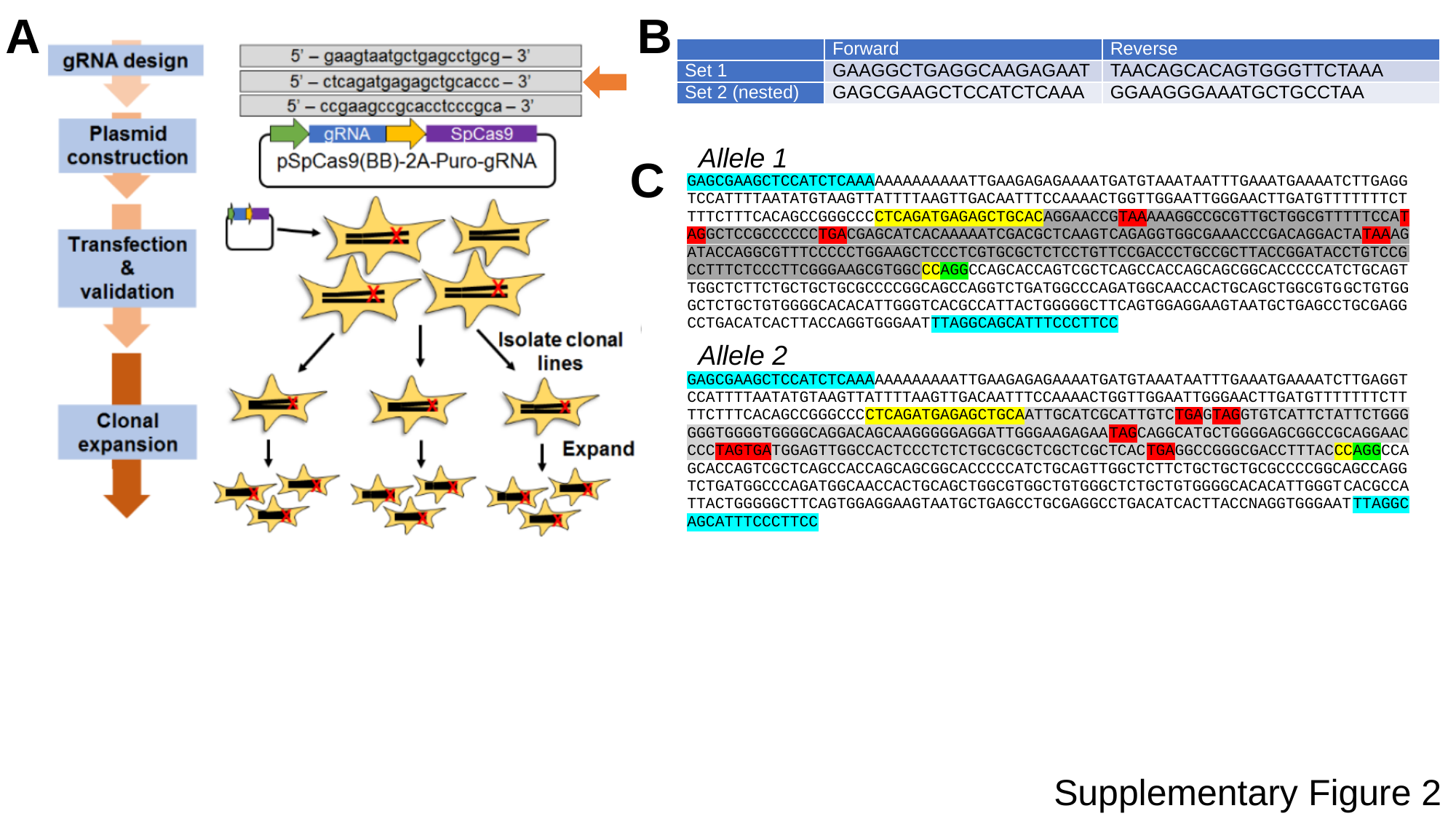

A
B
| | Forward | Reverse |
| --- | --- | --- |
| Set 1 | GAAGGCTGAGGCAAGAGAAT | TAACAGCACAGTGGGTTCTAAA |
| Set 2 (nested) | GAGCGAAGCTCCATCTCAAA | GGAAGGGAAATGCTGCCTAA |
Allele 1
Allele 2
C
Supplementary Figure 2
